# Opto-lipidomics of tissues

**DOI:** 10.1101/2023.04.01.534702

**Authors:** Magnus Jensen, Shiyue Liu, Elzbieta Stepula, Davide Martella, Anahid A. Birjandi, Keith Farrell-Dillon, Ka Lung Andrew Chan, Maddy Parsons, Ciro Chiappini, Sarah J. Chapple, Giovanni E. Mann, Tom Vercauteren, Vincenzo Abbate, Mads S. Bergholt

## Abstract

Lipid metabolism and signalling play pivotal functions in biology and disease development. Despite this, there is currently no optical technique available that can directly visualise the lipidome in tissues. In this study, we introduce opto-lipidomics, a new approach to optical molecular tissue imaging. We expand the capability of vibrational Raman spectroscopy to identify individual lipids in complex tissue matrices through correlation with desorption electrospray ionisation (DESI) - mass spectrometry imaging in an integrated instrument. A computational pipeline of inter-modality regression analysis is established to extract lipidomic information from optical vibrational spectra. Opto-lipidomic imaging of transient cerebral ischemia-reperfusion injury in a murine model of ischemic stroke demonstrates the visualisation and identification of lipids in disease with unprecedented molecular specificity using light. Furthermore, we deploy opto-lipidomics in a handheld fiber-optic Raman probe and demonstrate real-time classification of bulk brain tissues based on specific lipid abundances. Opto-lipidomics opens a host of opportunities to study lipid biomarkers for diagnostics, prognostics, and novel therapeutic targets.

## Introduction

Lipids, which include triglycerides, phospholipids, and sterols, are essential molecules providing fundamental building blocks of cells and tissues. Lipids are key components of plasma membranes, and the makeup of these membranes affects many biological functions, such as enzyme regulation and receptor activity. Additionally, certain types of lipid structures, such as lipid rafts, have crucial roles in maintaining proper cellular activity and function. Many diseases, such as ischemic stroke, atherosclerosis, neurological diseases, and cancers, are all associated with dysregulation in lipid content^1 2^.

Although lipids play a crucial role in many biological processes, imaging strategies to investigate them are limited since they are not amenable to labelling with fluorophores. To perform lipidomic analyses, scientists currently depend heavily on high-performance liquid chromatography (HPLC) and mass spectrometry (MS) of bulk tissues^3^. Spatial omics and imaging approaches provide unprecedent opportunity to study how information at the molecular scale contribute and shape tissue functions and phenotype. For instance, desorption electrospray ionisation (DESI)-MS has emerged as a powerful approach for conducting spatial lipidomics of tissue samples^4-7^. Optical vibrational techniques such as Fourier transform - infrared (FT-IR) or Raman spectroscopy can also be used to measure lipids in tissues^8^. Raman spectroscopy uses laser light to probe specific vibrational modes associated with the structure of molecules^9^. Raman spectroscopy holds great appeal for tissue analysis both *ex vivo* and *in vivo* due to its non-destructive nature, label-free detection scheme, and minimal sample preparation requirements^10,11^.

Optical spectroscopy and MS generally provide complementary information about tissue composition^12-15^. DESI-MS – typically involving high-resolution MS, offers precise and untargeted detection and identification of lipids and metabolites based on mass to charge ratios and determination of elemental composition. Raman spectroscopy probes the specific molecular vibrations in proteins, lipids, carbohydrates and DNA, enabling the analysis of subtle structural and conformational molecular differences^8^. While Raman spectroscopy can uncover the overall lipid content, it remains extremely difficult to distinguish different lipid subtypes in tissue matrices. This is due to the complexity caused by overlapping Raman peaks from the myriad of molecules in tissues resulting in a compound tissue spectrum^16^. This represents a monumental challenge for any vibrational spectroscopy technique in biomedical sciences. Raman spectroscopy has, however, unexploited potential to offer massively multiplexed tissue analytics *ex vivo* and *in vivo*. Existing methods for analysing compound tissue spectra rely predominantly on multivariate regression analysis using libraries of purified chemicals, principal component analysis (PCA)^17-19^ and clustering of molecular fingerprints^20^. While these computational techniques are efficient for estimating the overall chemical composition (e.g., total protein, lipid, and DNA content), in most cases they do not facilitate accurate identification and subtyping of lipids in tissues^21,22^. Sequential imaging using MS and Raman spectroscopy has therefore been reported for tissue analysis^12-14,23-26^. The utilisation of Raman and MS information enhances tissue characterisation owing to their complementarity.

Here we introduce a new strategy for optical lipid imaging. We expand the capability of vibrational Raman spectroscopy to identify individual lipids in complex tissue matrices. Development of the first integrated Raman and DESI-MS instrument enabled us to perform fully correlative lipid imaging of tissues. This method, “opto-lipidomics”, allowed us to develop a regression model for inferring lipidomic information from optical vibrational spectra. We demonstrate the use of opto-lipidomics by imaging ischaemia-reperfusion brain injury in a murine model of transient ischemic stroke showing that Raman spectroscopy can detect and distinguish specific lipids in complex tissues. Finally, we deploy opto-lipidomics in a handheld fibre-optic Raman probe demonstrating real-time bulk tissue lipidomics. Our method achieves for the first time direct optical visualisation and detection of specific lipid sub-types in complex tissues promising to catalyse lipidomic analysis *ex vivo* and *in vivo*.

## Results

### Integrated Raman/DESI-MS imaging

Sequential optical spectroscopy and MS imaging present significant technical hurdles, including the need for precise resolution matching, absolute pixel-to-pixel alignment, diverse sample preparation methodologies, and requirement of fast imaging to avoid degradation and evaporation of volatile compounds from the tissue. Moreover, correlating data from multiple modalities necessitates sophisticated computational tools. Raman spectroscopy and DESI-MS imaging exhibit remarkable compatibility, as they both employ label-free detection schemes. Integrating these modalities into a single instrument mandates a thorough evaluation of their compatibility. We developed an integrated Raman/DESI-MS instrument with matched imaging resolution (Fig. 1a-c) (*see Material and Methods*). The spatial resolution for correlative imaging is limited to ∼50 µm by DESI-MSI. Raster scanning is used to acquire data across spatial regions of interest and is controlled using an in-house developed software suite to enable electronic synchronisation with the motorised stage (Supplementary Fig. 1). Critically, we verified that the electrically charged MeOH jet spray did not give rise to interfering Raman peaks for any flow rates (Supplementary Fig. 2). This is due to the comparatively low detection sensitivity of Raman spectroscopy. We created protocols for Raman/DESI-MS imaging alignment and substrate preparation (*see Material and Methods*). Since Raman spectra are sampled at an angle we characterised and found only subtle angular dependence (Supplementary Fig. 3). Characterisation of the impact of tissue thickness on the Raman signal generation indicated that 40 µm (corresponding to ∼2-3x of cell size) provided a reasonable compromise between tissue thickness and signal intensity generation (Supplementary Fig. 4). DESI-MS imaging is a partially destructive technique since the tissue is exposed to MeOH and desorption/ionisation of the superficial tissue layer occurs (Supplementary Fig. 5). We, therefore, investigated the impact of DESI-MS imaging on the Raman signal, but we found no significant differences induced by DESI-MS as revealed by PCA scores (Supplementary Fig. 6). Integrated Raman and DESI-MS imaging generate vast amounts of hyperspectral data that require independent preprocessing. We created a computational pipeline for processing the two modalities (Fig. 1d), including background image masking, separate preprocessing for Raman/MS (Supplementary Fig. 7), pixelwise co-registration and heterospectral correlation analysis (*see Material and Methods*). Finally, we investigated the complementarity between optical spectroscopy and DESI-MS by measuring structural isomers bearing identical elemental composition that cannot be differentiated using DESI-MS alone (Supplementary Fig. 8). These results demonstrates that while DESI-MS can offer specific identification of the lipid sub-class, Raman spectroscopy can probe the subtle vibrational differences reflecting the molecular structure and resolve highly similar lipid species.

**Fig. 1.**
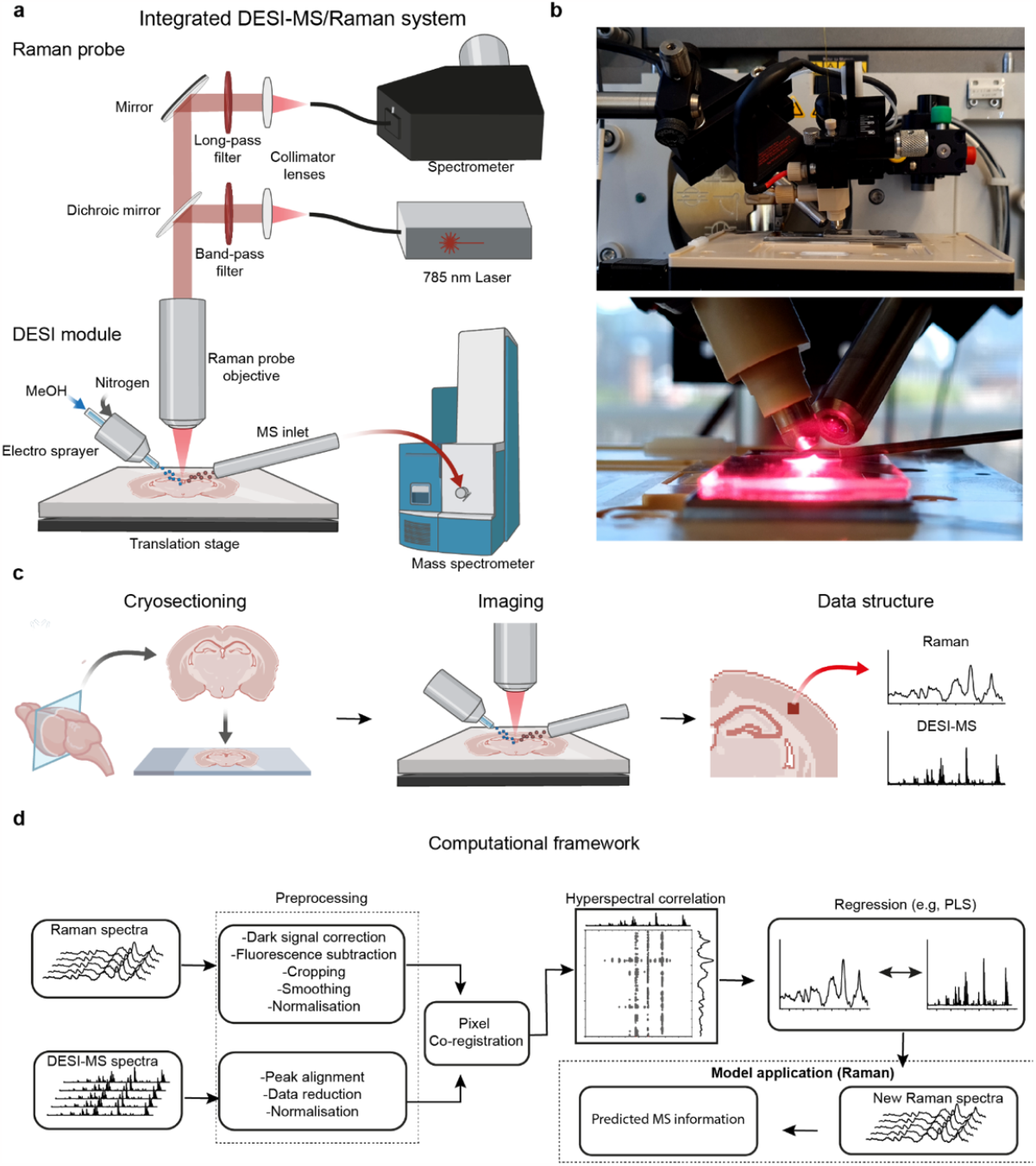
Integrated Raman and DESI-MS imaging system. **a**, Schematics of the integrated Raman and DESI-MSI system. The electrosprayer pneumatically focuses a jet of electrically charged MeOH solvent on the tissue sample using pressurised nitrogen. Desorbed molecules are then passed through ambient air into the mass spectrometer (MS) inlet. For Raman spectroscopy, a 785 nm laser light is delivered through an objective lens and focused onto the sample at a 45-degree angle. Raman scattered is passed back through the objective and separated from the laser path by a dichroic mirror and longpass filter before it is fiber-coupled into a near infrared (NIR) spectrometer. **b**, Photograph showing the integrated Raman/DESI-MS imaging system using a commercial Raman probe (InPhotonics). **c**, Integrated Raman/DESI-MS imaging workflow for tissue analysis including cryosectioning, correlative imaging and analysis. **d**, Computational framework for analysis: Data processing and correlation analysis pipeline for Raman and DESI-MS data. The Raman and DESI data are preprocessed separately before they are co-registered. Heterospectral correlation is performed to ensure accurate co-registration and can be used to assess complementarity and interrelationship. Lastly, a regression model can be constructed to predict relative m/z abundances from the Raman spectra.

### Opto-lipidomic imaging of mouse brain tissues

Using the integrated Raman/DESI-MS imaging instrument, we imaged 8 sections from brains of healthy mice as these exhibit well-defined anatomical landmarks of white and grey matter (n = 384,000 Raman spectra and n = 384,000 DESI-MS spectra) (Fig. 2a-d). DESI-MSI false-colour images allow visualisation of the presence of prominent lipids and their relative abundances: putatively PI 38:4 (m/z 885.6), ST 24:1 (m/z 888.6), PS 40:6 (m/z 834.5) associated with white and grey matter (Fig. 2a, Supplementary Fig 8-9). The mean DESI-MS spectrum for the entire tissue sample showed distinct m/z peaks previously reported (Fig. 2b)^27^. Correlation analysis revealed that the samples contained a significant number of lipid species that were independently distributed in the tissue (Supplementary Fig. 10). In contrast to DESI-MS, which exhibits high molecular specificity, the average Raman spectra of tissues contained numerous overlapping vibrational peaks, resulting in limited molecular specificity (Fig. 2c-d). However, by imaging prominent Raman peaks, such as 1445 cm^-1^ (CH_2_ deformation of lipids), 1650 cm^-1^ (υ(C=C) of lipids + Amide I of proteins) and 1060 cm^-1^ (tentatively assigned to DNA), we observed similar anatomical landmarks.

**Fig. 2.**
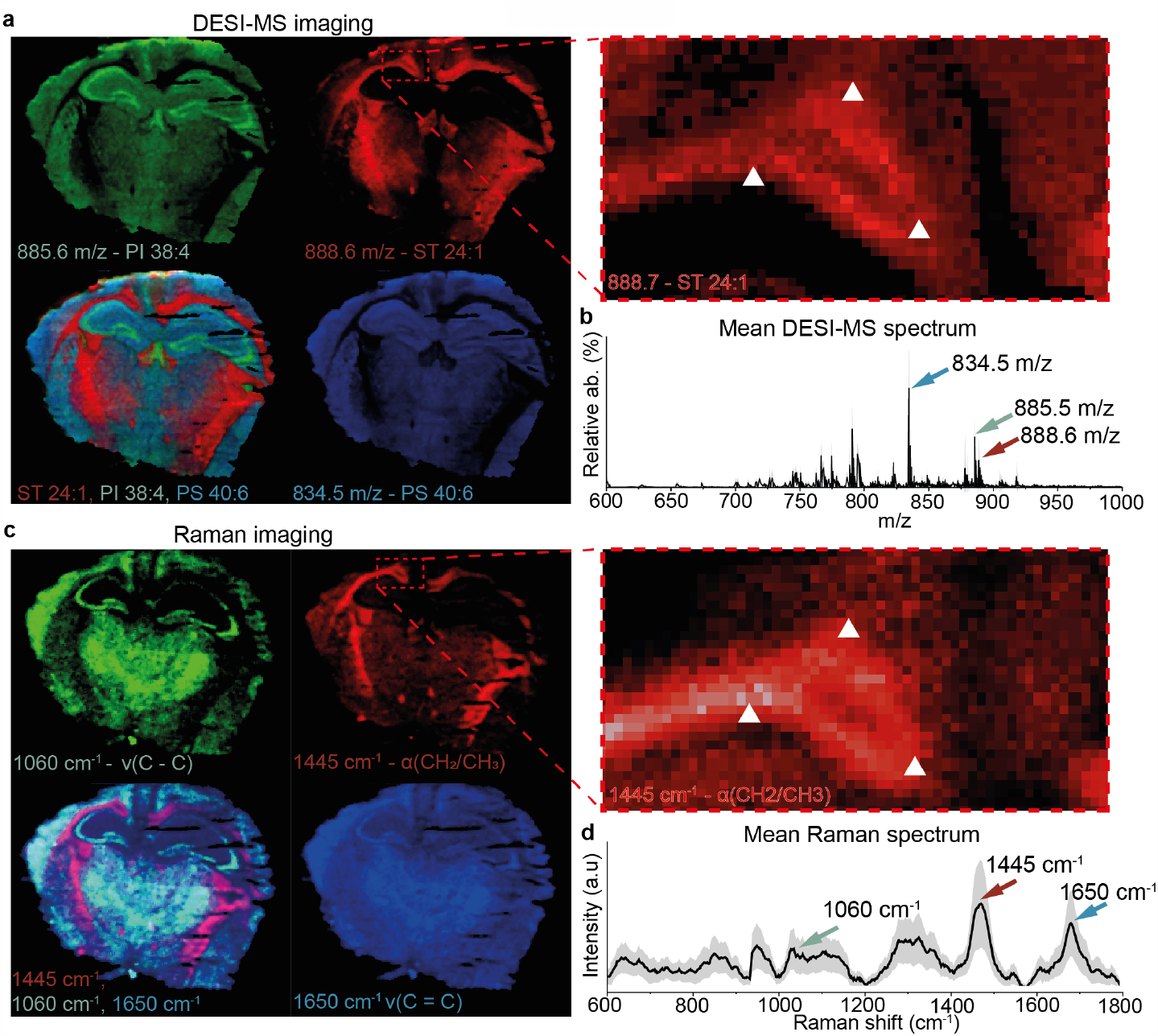
Co-registered Raman/DESI-MS imaging of healthy mouse brain tissue. Co-registration of the Raman and DESI-MS hyperspectral images both sampled at 50 µm resolution. **a**, DESI-MS image of ST 42:1, PI 38:4 and PS 40:6. White triangles in Raman and DESI-MS images indicate markers of similar anatomical location. **b**, Mean DESI-MS spectrum of brain tissue. **c**, Raman images generated from peaks at 1445 cm^-1^, 1060 cm^-1^, and 1650 cm^-1^ tentatively assigned to CH_2,_ ν(C-C), and ν(C=C) respectively. White triangles in Raman and DESI-MS images indicate markers of similar anatomical location. **d**, Mean Raman spectrum ±1 standard deviation (SD) of all tissue spectra. Triangles in Raman and DESI-MS images indicate markers of similar anatomical location.

Pixel-wise correlation analysis (*see Materials and Methods*) demonstrated complex yet interpretable relationship between tissue Raman peaks and MS lipid abundances (Fig. 3a). For example, the strong correlation between the 1445 cm^-1^ Raman peak (CH_2_ deformation of lipids) and m/z 888.6 verified the Raman contrast of white matter.

**Fig. 3.**
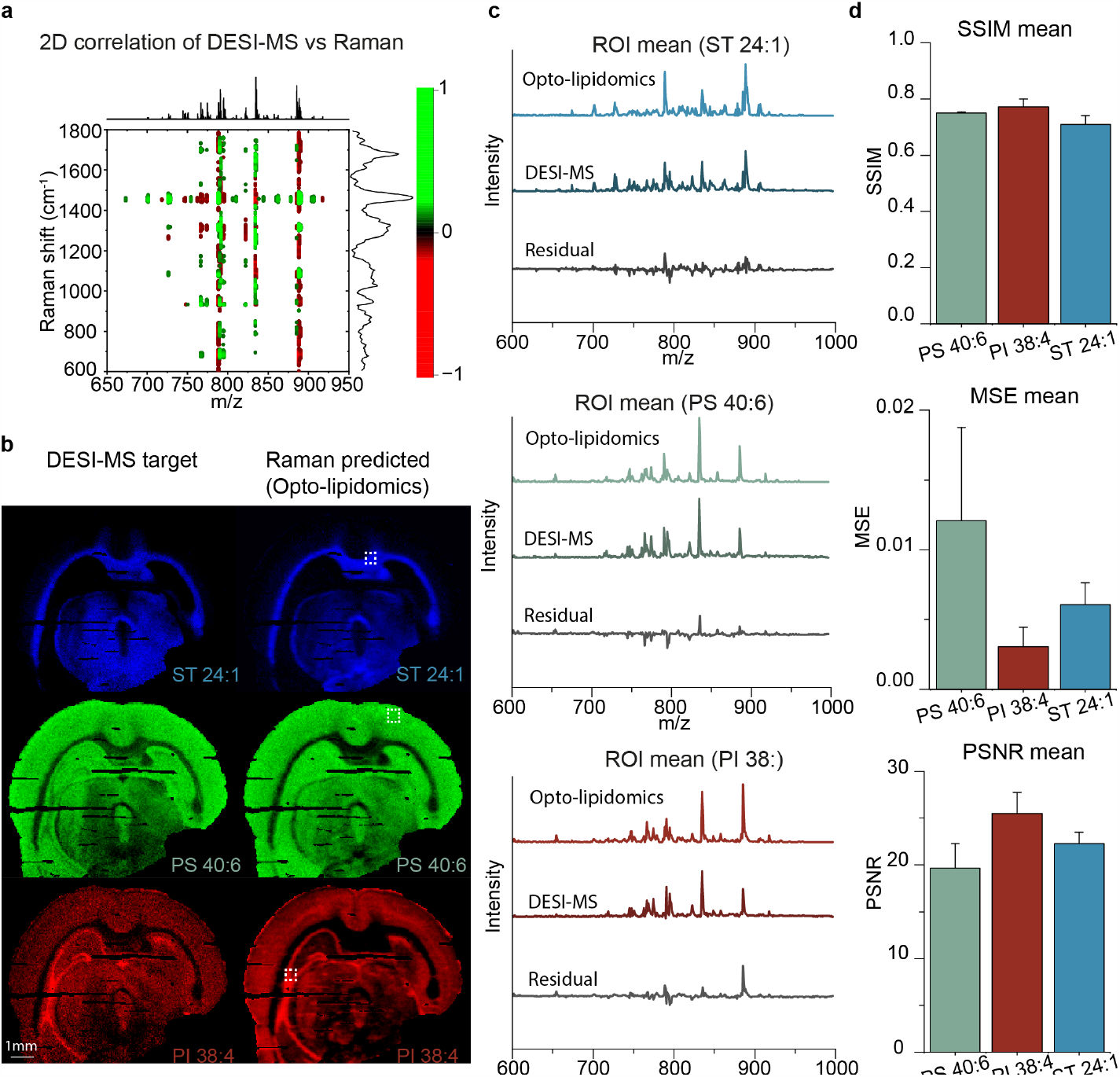
Opto-lipidomic imaging of healthy mouse brain tissue. **a**, 2d heterospectral correlation between Raman and DESI-MS for healthy mouse brain tissue. **b**, Images of a coronal mouse brain section showing target DESI-MS and Raman predicted (opto-lipidomics) images: ST 24:1, PS 40:6, and PI 38:4. **c**, Representative mean spectra of region of interest (ROI)s associated with the three lipids using DESI-MSI and Raman predicted (opto-lipidomics). Also shown are the residual spectra. **d**, Image quality metrics: structural similarity index (SSIM), mean squared error (MSE) and peak signal to noise ratio (PSNR), for PS 40:6, PI 38:4, and ST 24:1 (n=3) (error bars: mean ± standard deviation (SD)).

Our goal was to infer lipidomics from optical vibrational spectra. We, therefore, performed inter-modality partial least squares (PLS) regression analysis to predict the m/z abundances from the Raman spectra. The developed regression model was used to predict the m/z abundances in 3 sections from one independent mouse brain (n=70,389 Raman spectra and n=70,389 DESI-MS spectra). We imaged the distribution of three of the most abundant m/z peaks as well as the Raman predicted (opto-lipidomic) distributions (Fig. 3b). The Raman predicted lipid distributions images showed striking similarities with DESI-MSI such as nearly indistinguishable distribution of PS 40:6 in the cerebral cortex and brain stem. The average DESI-MS spectrum and Raman predicted (opto-lipidomic) spectrum for the three regions of interest (ROI) also showed good correlation (Fig. 3c). The residual difference spectrum reveals that the model showed poor performance for some m/z peaks (e.g., m/z 766.6, m/z and 794.6). The images of the 12 most prominent DESI-MS peaks and their predicted lipid distributions can be found in Supplementary Fig. 11. We quantitatively assessed the degree of confidence in the Raman prediction of the abundance of the respective lipids by calculating the structural similarity index (SSIM)^28^, mean square error (MSE) and peak signal-to-noise ratio (PSNR) (Fig. 3d). The congruent distribution of lipids revealed by opto-lipidomics and DESI-MS established the feasibility of using MS to infer omics information from optical vibrational spectra.

### Opto-lipidomic imaging in a murine model of transient cerebral ischemia-reperfusion injury

We next investigated if opto-lipidomic imaging could be used to identify alterations in lipids in a disease model. Ischemic stroke is a leading cause of mortality and adult disability, and restoration of cerebral blood flow is currently the only effective treatment. Cerebral ischemia/reperfusion injury is associated with increased generation of reactive oxygen species^29^, mitochondrial dysfunction^30 31^, and dysregulation of lipid metabolism^32^, yet studies of altered lipid metabolism in ischemic stroke are limited. We imaged 3 tissue sections from a mouse model of transient cerebral ischemia-reperfusion injury (n=77,032 Raman spectra and n=77,032 DESI-MS spectra) whereby a one-sided reperfusion injury is induced while the contralateral brain hemisphere represents internal control tissue. Regression analysis was used predict the m/z abundances in an independent mouse brain section with one-sided reperfusion injury (Fig. 4a-c). These results show that opto-lipidomic imaging largely replicated the DESI-MSI results. Region-of-interests (ROIs) in injury and control side (Fig. 4b-c) revealed that FA 22:6 (m/z 327.2 associated with docosahexaenoic acid (DHA)) and CER 36:1;O2 (m/z 600.5) were accumulated in the injured hemisphere. Most importantly, these Raman-derived observations from the ROI agreed with the conclusions drawn from DESI-MSI (Fig. 4d-e-f). Image quality metrics (SSIM, PSNR and MSE) revealed that the predicted lipid distributions recapitulate the DESI-MSI results (Supplementary Fig. 12). This data indicates that opto-lipidomics can provide a reliable imaging strategy for providing molecular insights into diseases.

**Fig. 4.**
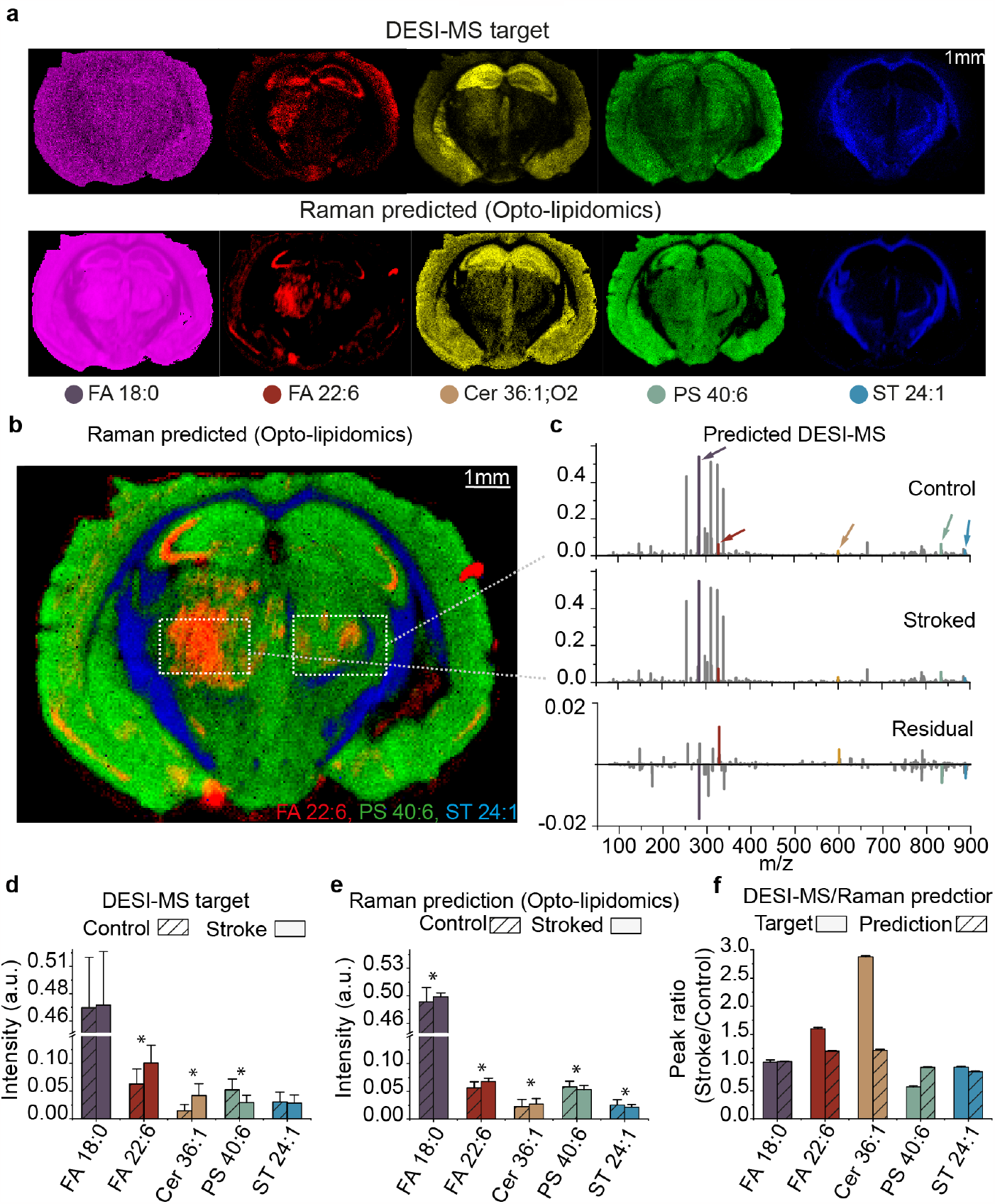
Opto-lipidomics in a mouse model of transient cerebral ischemia-reperfusion injury. **a**, Images of a coronal mouse brain section post ischemia-reperfusion showing DESI-MSI target and Raman predicted (opto-lipidomic) images associated with the brain at FA 17:1, FA 22:6, Cer 36:1;O2, PS 40:6 and ST 24:1. **b**, Merged Raman predicted (opto-lipidomic) distribution of FA 22:6, PS 40:6, and ST 24:1. **c**, Representative mean predicted MS spectra for the stroked and contralateral control side. Also shown is the residual. **d**, DESI-MS target peak intensity values in stroked and control side ROI for FA 17:1, FA 22:6, Cer 36:1;O2, PS 40:6, and ST 24:1. **e**. Raman predicted peak intensity values in stroked and normal side ROI for FA 17:1, FA 22:6, Cer 36:1;O2, PS 40:6, and ST 24:1. **f**, DESI-MS target and Raman predicted peak residual values between stroked and normal side ROI for FA 18:0, FA 22:6, Cer 36:1;O2, PS 40:6, and ST 24:1. (Data denote mean ± standard deviation (SD), * = Significant difference between stroked and control side, P<0.05).

### Handheld real-time opto-lipidomics of bulk tissues

As a final demonstration of opto-lipidomics, we obtained an intact mouse brain and excised it in the transverse plane (Fig. 5a). We then used a handheld fibre-optic Raman probe for real-time (0.5 sec acquisition) sampling across the white and grey matter (Fig. 5b). The regression model previously trained on (n = 130,773 Raman and n = 130,773 DESI-MS spectra) was used to predict the m/z abundances in bulk tissue. We acquired Raman spectra of normal mouse brain containing visually apparent white matter (n=10) and grey matter (n = 10) (Fig. 5c). Due to diffuse light scattering in bulk tissues, there were only subtle differences in the Raman spectra of white and grey matter. Nevertheless, opto-lipidomic predictions of lipid abundances (ST 24:1 vs PS 36:1, ST 24:1 vs PI 38:4 and ST 24:1 vs PS 40:6) still demonstrated 100% discrimination between white and grey matter (Fig. 5e). These results highlight the first proof of concept of a workflow for opto-lipidomics of bulk tissues.

**Fig. 5.**
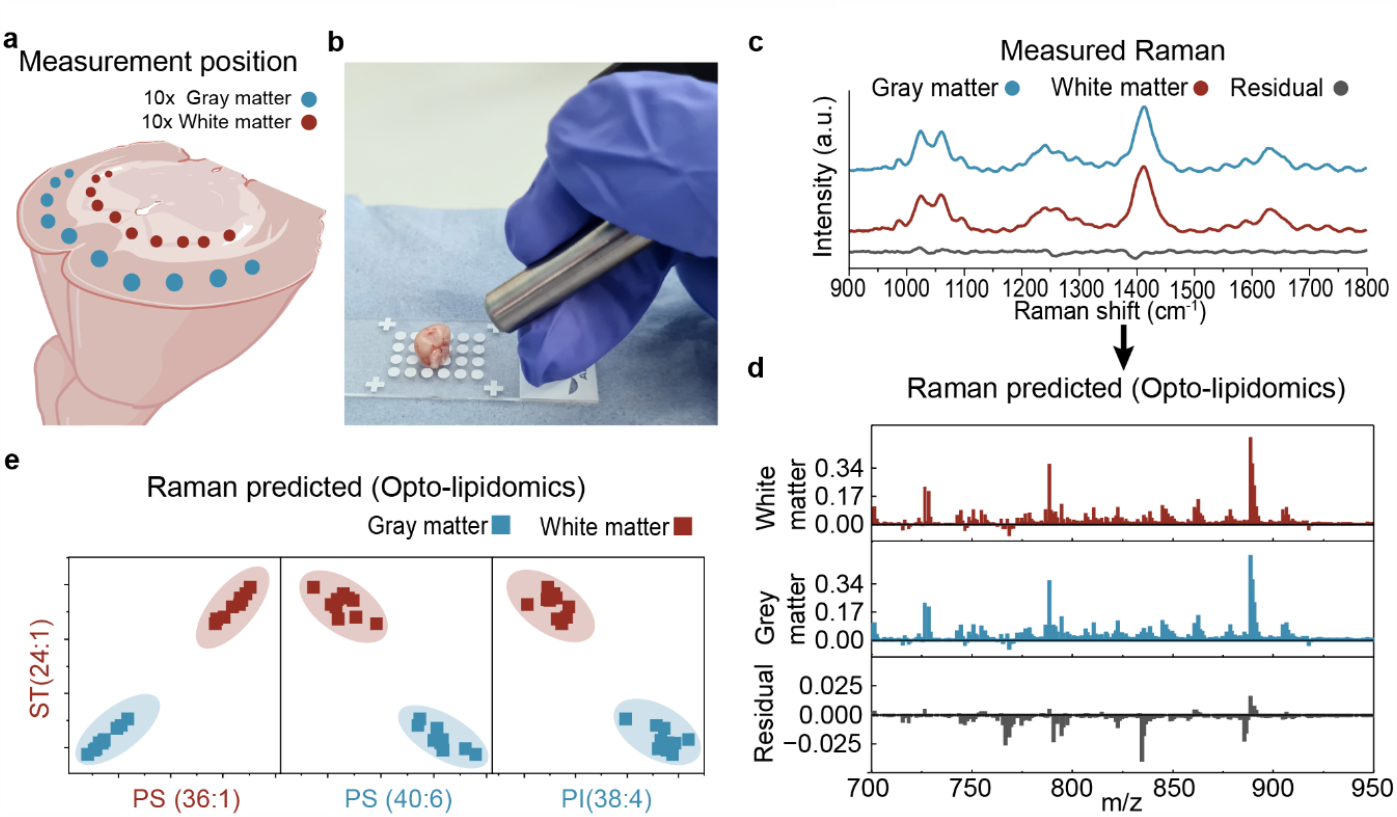
Handheld real-time opto-lipidomics of bulk tissues. **a**, Schematic of coronal cut mouse brain with measurement position represented in green (grey matter) and blue (white matter). **b**, Image of handheld Raman probe and coronal cut mouse brain. **c**, Representative mean Raman spectra for the grey matter, white matter as well as the residual difference spectrum. **d**, Representative mean predicted DESI-MS spectra for the grey matter, white matter. Also shown is the residual. **e**. Scatter plot depicting the predicted content of PS 36:1, PI 38:4, PS 40:6 against ST 24:1, showing separation of white and grey matter measurements (n=20).

## Discussion

Currently, there is a lack of optical microscopy tools that can directly visualise the lipidome in tissues. However, we present a new approach that can generate optical microscopic contrast of lipids. While Raman spectroscopy is sensitive to lipids, its ability to accurately identify lipid species within complex tissue matrices is inherently limited. We have demonstrated that Raman spectroscopy and DESI-MS exhibit complementarity in their ability to analyse lipids in a controlled experiment but also that they can be sensitive to similar molecules. We showed that this can be leveraged for lipidomic inference of Raman spectra. We engineered a fully integrated Raman and DESI-MS imaging instrument enabling us to increase throughput and enable precise co-registration.

We demonstrated the application of opto-lipidomics using healthy and diseased mouse brain tissue because the anatomical landmarks are well-defined. For many of the most abundant DESI-MS peaks, the Raman prediction and image reconstruction performed exceptionally well in replicating the DESI-MS m/z abundances; however, for some lower-abundance DESI-MS peaks, the predictive capability was reduced (Supplementary Fig. 9). This is expected and can be attributed to a number of limitations, including the low detection sensitivity of Raman spectroscopy, the presence of MS fragments/adducts and isomers, as well as the general inadequacy of linear regression modelling. Application of opto-lipidomics to a murine model of transient cerebral ischemia-reperfusion injury revealed a significant increase of FA 22:6 (docosahexaenoic, DHA) in the stroke hemisphere that has previously been associated with neural protection^33,34^. Docosanoids have been shown to increase neurogenesis, angiogenesis and, importantly, synthesis of one of its derivatives, neuroprotectin D1 in the penumbra of the stroke hemisphere, thereby potentially affording protection against further ischemic damage. Moreover, pro-inflammatory ceramide 36:1;O2 (CER) was also shown to accumulate within the stroked hemisphere, with previous studies showing a correlation between plasma CER accumulation and stroke risk and severity^35,36^. These findings showcase the utility of opto-lipidomics in investigating lipid biomarkers for applications in studying disease and potential identification of therapeutic targets^37,38^.

Handheld opto-lipidomics of bulk tissues serve as an important proof of concept of how lipidomic information can be extracted in real-time using light. This is in sharp contrast to state-of-art computational methods for Raman spectroscopy, such as PCA, which does not offer molecular identification. Since Raman spectroscopy can be performed in humans^39,40^, the developed approach paves the way for *in vivo* optical lipidomics. *In vivo* applications, on the other hand, would necessitate rigorous validation (i.e., due to hydration/vascularity) such as potential correlation with bulk tissue LC-MS analysis.

There are many prospects for improving the predictive capability of our imaging approach. Due to various experimental factors, such as total laser power, sample dimensions, electrospray flow rate, and the size of the electro-sprayer reservoir, the minimum integration time per pixel was determined to be approximately ∼0.5 seconds. Increasing integration time might improve the detection limit of Raman spectroscopy while resulting in extended imaging time. Furthermore, here we employ PLS regression since it offers straightforward model interpretability. Since PLS regression relies on linear data projections, we anticipate that utilising more advanced non-linear computational methods would enhance the limit of detection and molecular specificity. DESI-MS examines only a shallow layer of the tissue, whereas Raman spectroscopy with NIR laser excitation probes the entire 40 µm tissue thickness. Therefore, developing specialised focusing optics with extended working distance and genuine confocal capability, or even visible laser excitation, could potentially enhance the correlation even more.

The imaging methodology is subject to important limitations. Spectral normalisation is necessary for both Raman and DESI-MS imaging to enable comparative analysis between images. As a result, the regression provides solely a relative molecular analysis. Importantly, if the tissue is highly homogenous or if multiple lipid species covariates, this can lead to deceitful regression analysis. However, we anticipate that with the most recent development of DESI-MS sprayers with imaging resolution down to 8-20 micrometres this will allow us to uncover much more details of the extracellular matrix and even offer single-cell opto-lipidomics which will likely improve analysis of more homogenous tissues. Interestingly, recent development in DESI-MS has shown the capability of imaging proteins and peptides using tissue digestion techniques which will massively increase the scope of this methodology^41^. Lastly, it should be noted that the regression model was developed for a particular tissue type and Raman system and thus need to be validated prior to implementation for other issues. Our future objective is to introduce deep learning-based analysis and transfer learning approaches, which will facilitate the expansion of the modelling capability to additional tissues and systems with minimal effort^42,43^. Potential pragmatic advancements could involve incorporating additional modalities, such as FTIR, into the Raman/DESI-MS system. Exploring directions for functional imaging at the molecular, cellular, and tissue levels may entail correlating higher resolution opto-lipidomics with complementary analyses such as proteomics and transcriptomics.

In summary, we developed a unique integrated Raman and DESI-MS imaging instrument. This enabled us to expand the capability of vibrational Raman spectroscopy to identify individual lipids in tissues. This creates novel possibilities for tissue characterisation and highlights a new strategy for implementing optical techniques in the imaging and examination of lipid biomarkers in diseases.

## Methods

### Development of integrated Raman and DESI-MSI instrument

For DESI-MSI, an electrosprayer pneumatically focuses a jet of electrically charged MeOH solvent onto the tissue sample using pressurised nitrogen. The DESI-MS system (Waters Corporation) comprises a DESI module interfaced to a high-resolution hybrid quadrupole-time of flight mass spectrometer (XEVO G2-XS QTOF MS) operated in sensitivity mode (Negative ion, MS resolution: 2000, Window: 0.02). The DESI module encompasses a sprayer head and a motorised sample stage. Nebulising nitrogen gas is externally supplied to the system. An electrospray solvent (HPLC-grade MeOH, ≥99.9%, Sigma Aldrich67-65-1) is passed to the electro-sprayer head through a fused silica capillary tube (Waters 6490512-S3) attached to a 10 ml syringe (SGE Model 1MR-GT 1 ml/23/2). The syringe is placed into a syringe pump (Harvard Apparatus, Pump 11 Elite), which applies pressure to the syringe, slowly supplying a constant flow of solvent to the sprayer head. The sample stage controller is connected to the mass spectrometer controlled by a workstation. The imaging resolution of the DESI-MSI system is ∼50 µm with a spectral resolution of 0.02 m/z. We implemented an in-house customised near-infrared (NIR) Raman modality with long working distance objective as well as an interchangeable commercial probe (InPhotonics Inc) (Fig 1a). At a 45-degree angle, we focused 785 nm laser light onto the same spot as the electrosprayer to enable correlative Raman spectroscopy. The Raman scattered light was passed back through the collection optics and focused onto a 100 µm fibre acting as an optical pinhole before entering a high-throughput NIR-optimised spectrometer. For all imaging here, we employed a commercial NIR Raman probe (Ocean Insight RPB785) with backscattering geometry in the DESI module. The Raman probe was connected to a 785 nm laser (B&W Tek BRM-785-0.55-1000.22-FC, 600 mW) with a 105 µm excitation fibre and to a NIR spectrometer (Princeton Instruments Acton LS785 using a Princeton Instruments PIXIS 400BR 750–1100 nm) with a 200 µm collection fibre. A digital acquisition board (National Instruments USB X SERIES Multifunction DAQ) was connected to the I/O control box of the DESI-MS system and used to synchronise the acquisition.

### Fibre-optic Raman system for bulk tissue spectroscopy

We utilised a NIR Raman probe (Ocean Insight RPB785) to perform the bulk tissue measurements. The Raman probe was connected to a 785 nm laser (B&W Tek BRM-785-0.55-1000.22-FC, 600 mW) with a 105 µm excitation fibre and a NIR spectrometer (Princeton Instruments Acton LS785 using a Princeton Instruments PIXIS 400BR 750–1100 nm). A comprehensive software has been developed in the Matlab 2022a environment to provide real-time data collection and analysis.

### Raman/DESI-MS imaging alignment protocol

To align the two modalities at the micron level, we have created an alignment protocol using lines of calibration ink on magnesium fluoride (MgF_2_) substrate that gives rise to intense fluorescence and DESI-MS signal. We iteratively optimised the alignment of Raman spectroscopy and DESI-MS in the x-y lateral plane to achieve maximum fluorescence and MS signal. First, the DESI-MS sprayer was focused and optimised on a magnesium fluoride (MgF_2_) slide (Global Optics Ltd) with black calibration ink. While synchronously measuring the signal intensity of the black dye on both the DESI-MS and Raman systems (fluorescence), the Raman focusing module was optimized in the x-y plane for maximum intensity. This was performed multiple times in both X and Y directions until the focusing spots of the two systems overlap.

### Raman/DESI-MS imaging of brain tissues

All Raman and DESI-MS spectra were collected on the integrated Raman/DESI-MS system simultaneously. Coronal cut mouse brain sections were imaged (∼10000 × 12000 µm (spatial resolution of 50 µm)) by raster scanning (Supplementary Fig. 1) across the entire tissue surface. Each Raman and DESI-MS spectrum was collected with an acquisition time of ∼0.5 s and laser power on the sample of 200 mW which did not give rise to sample degradation due to relatively low laser power density. The electrosprayer supplied MeOH at a flow rate of 2 µl/min, with a nebulising gas (nitrogen) pressure of 0.5 bar. The electrosprayer was angled at 65 degrees, and the Raman probe at 40 degrees to the surface of the tissue.

### Charged MgF_2_ substrate

Raman spectroscopy is poorly compatible with standard microscope glass slides and requires Raman-compatible substrates such as MgF_2_ or Raman grade CaF_2_. For this reason, we developed a protocol to electrostatically charge MgF_2_ slides using poly-l-lysine, making Raman spectroscopy and DESI-MS imaging fully compatible for correlative tissue imaging. This is necessary since the DESI gas pressure can destroy tissue that is not adhering to the substrate. 1% poly-l-lysine in H_2_O (Merck SLCL3300) was mixed with milli-Q water in a ratio of 1/9. Cleaned MgF_2_ slides were then placed into a petri dish, and the poly-l-lysine solution was poured over the slides leaving ∼10 mm of the solution above the slides. The slides were then stored in the solution for 24 hours at 25°C, with a cover on top to avoid contaminants. Afterwards, the slides were removed from the solution, air-dried, and stored in a microscope slide container at room temperature.

### Healthy mouse brain sample preparation

Female Albino CD1 wildtype mice aged 6 weeks were used in the experiments in accordance with UK Home Office Procedures. Brain dislocations were carried out immediately after animal sacrifice by neck dislocation. Using sterile instruments, the skull was exposed by cutting the skin on top of the head. Occipital and interparietal bones were cut. An incision was then made in the skull along the sagittal and parietal sutures. A hole was made in the skull at the junction of frontal and parietal bones, where the tip of a scissor was inserted to crack open the calvaria. The remaining skull was removed to expose the brain completely and remove the remaining nerves and peduncles. The brain was then taken out of the skull and immediately transferred to liquid nitrogen. The brains were stored at -80°C until sectioning. Coronal sections of the mouse brains were cryosectioned (Bright OFT Refrigerated Cryostat) with a 40 µm thickness and thaw mounted on charged MgF_2_ slides. The samples were stored at -80°C until analysis. Prior to imaging, the tissues were thawed at room temperature.

### Mouse model of transient cerebral ischemia-reperfusion injury

Male C57BL/6J mice, aged 10-12 weeks and maintained on a 12-h light cycle from 0700 to 1900, were used for experiments in accordance with UK Home Office Procedures. Mice were pre-treated *i*.*p*. with 5 mg/kg of methadone prior to the onset of surgery and induced to a surgical plane of anaesthesia with inhalation of 4% isoflurane in 30% oxygen, reduced to 1.0-1.5% isoflurane for maintenance. The middle cerebral artery was transiently occluded for 60 min, using a modification of the Longa technique^44^, followed by recovery from anaesthesia and reperfusion for 72 h with analgesia provided. Brains were washed in heparin/saline and directly flash frozen. Brains were cryosectioned with three sections (40 µm) cut near -2 mm relative to bregma, placed on electrostatically charged MgF_2_ slides and stored at -80°C until imaging.

### Purified lipid sample preparation and measurements

Two lipids (18:1-18:0 PC, 18:0-18:1 PC) (Merck) and stored at -20°C until sample preparation. A ratio of 2 mg/ml lipids to chloroform solution was prepared for each different lipid. Multiple droplets of the solutions were deposited separately on CaF_2_ slide discs. After air-drying for 10 minutes, a thin lipid film formed on the slide. The slides were then analysed on the DESI-MS system. For the Raman analysis, 5 mg of the dry lipids were placed on an MgF_2_ slide and measured using 785 nm laser excitation. The excitation laser power for Raman imaging was 74 mW under 50X dry objective lens with 1 second integration time. We report the average of n=20 spectra sampled at random locations for each lipid.

### MS/MS

Utilizing tandem mass spectrometry (MS/MS) fragmentation, species identification within mouse brain specimens was conducted via MS/MS. The electrospray ionization voltage was fixed at 4.5 kV, for the acquisition of negatively charged ions. Methanol was delivered at a flow rate of 5 μL/min and subsequently atomized through the application of nitrogen gas at a pressure of 5 bar. To determine the identities of MS/MS fragment ions, the fragmentation patterns of chosen mass spectrometry peaks were compared with those available in online MS/MS spectral repositories, specifically MassBank and Lipid Maps.

### Raman spectral preprocessing

The raw Raman spectra were first calibrated to reduce etalon artefacts (Supplementary Figure 7). The mean spectrum using an average of multiple reference measurements (n=10) was fitted with a 5^th^-order polynomial fit and subsequently divided with a polynomial fit of green glass fluorescence. This calibration was then multiplied onto all the raw spectra to suppress etalon artefacts^9^. For each tissue section, a region of 10×10 pixels outside the tissue area was empirically chosen, and the mean spectra of the region were used as a background spectrum for each tissue sample, respectively. All spectra were then background subtracted. The Raman processing adhered to standard methods including autofluorescence removal, smoothing and normalisation. In this work, we found that a 2^nd^-order constrained polynomial fitted to the spectra efficiently could remove the autofluorescence background. Each Raman spectrum was normalised to the integrated area to ensure comparability between images and finally smoothed using a Savitzky-Golay filter (zeroth order, window: 3).

### DESI-MS spectral preprocessing

The 1000 most prominent peaks of the DESI-MS images were exported to a text file using HDI software (HDI version 1.5) and imported into Matlab 2022a. In Matlab, the DESI-MS peaks were sorted using the functions [*mspeaks* (conversion of MS data to a peak list), *mspalign* (alignment of MS spectra in peak list to a selected range reference peaks), *msppresample* (resample aligned MS spectra while preserving selected peaks)] in the Matlab Bioinformatics toolbox. The spectra were up-sampled to 20000 spectral points (to ensure a resolution of 0.02 m/z) from 600-1000 m/z (phospholipid region) for the normal mouse brain experiment. For brain sections from mice subjected to ischemia-reperfusion, the spectra were up-sampled to 57500 spectral points (to ensure a resolution of 0.02 m/z) between m/z 50 – 1200 (metabolites-fatty acids-phospholipids). The up-sampled MS spectra were then peak-aligned using the Matlab function *mspalign*. After peak alignment, to reduce the memory size for computations, the healthy mouse brain spectra were down-sampled to 485 spectral points, between 600-1000 m/z. For brain sections from mice subjected to ischemia-reperfusion, spectra were down-sampled to 987 spectral points between 50-1200 m/z, based on the 987 most prominent peaks allowing us to analyse to the low m/z region.

### Image masking and background removal

A mask was created for each tissue section to remove irrelevant background data (e.g., pixels from the MgF_2_ substrate) by taking the spectral sum of each pixel and removing any pixels under a lower threshold determined empirically for each image. This efficiently removes the substrate background.

### Co-registration

The preprocessed Raman and DESI-MS images were spatially aligned with a feature selection approach using functions (*cpselect*) from the Matlab Image Processing Toolbox. An in-house developed alignment algorithm was used to calculate any misalignment and adjusts the images accordingly. This approach is adopted to ensure that the images are linearly aligned by whole pixels.

### Heterospectral correlation analysis

Heterospectral two-dimensional correlation analysis (2D-COS)^11^ was performed on the preprocessed Raman and DESI-MS data. The synchronous correlation spectra were used to investigate the relationship between Raman and DESI-MS peaks.

### Partial least squares (PLS) regression modelling

Partial least squares (PLS) regression models were systematically trained for each m/z peak in the DESI-MS spectra. The spectral data was split into a training and independent test data set (i.e., independent mice brain). We performed PLS regression of the Raman spectra against the individual DESI-MS spectral peaks in the training data set. Mean centring was performed to reduce model complexity. We used an optimised number of latent variables (LVs) determined by leave-one-section-out cross-validation of the training set to minimise the root mean square error of cross-validation (RMSECV). The test data set was fully independent and used to validate the regression model by predicting the DESI-MS spectra from the corresponding Raman spectra. To quantify the prediction capabilities for each m/z abundance, we calculated the image quality metrics, PSNR and SSIM, as well as MSE. All computation in this work was performed on a workstation (AMD Ryzen) with 64 GB of memory.

### Statistical analysis

Multiple comparisons were calculated using one-way analysis of variance (ANOVA) in Origin Pro 2022b.

## Acknowledgements

This work has received funding from the European Research Council (ERC) under the European Union’s Horizon 2020 research and innovation programme (M.S.B., grant agreement No. 802778 and C.C. grant agreement 759577) and the Wellcome Leap Delta Tissue Programme. This work was also supported by the Engineering and Physical Sciences Research Council [M.S.B., grant number EP/S017607/1] and studies of ischemic stroke by the British Heart Foundation [FS/19/25/34277, G.E.M.].

## Contributions

M.J. performed the experiments, interpreted the data, generated the figures, and wrote the manuscript. S.L. performed experiments, contributed to scientific discussion, and data analysis. E.S., D.M., performed experimental work and contributed to scientific discussion. A.A.B, K.F.D, S.C., and G.E.M. contributed with scientific discussion, animal experiments and data interpretation. K.L.A.C, M.P, and C.C contributed to the scientific discussion and data interpretation. T.V. contributed with scientific discussion, data interpretation and data analysis. V.A contributed with scientific discussion, instrument development and MS data interpretation. M.S.B. designed the study, interpreted the data and wrote the manuscript.

## References

1 Santos, C. R. & Schulze, A. Lipid metabolism in cancer. FEBS J 279, 2610–2623 (2012). https://doi.org:10.1111/j.1742-4658.2012.08644.x

2 Natesan, V. & Kim, S. J. Lipid Metabolism, Disorders and Therapeutic Drugs - Review. Biomol Ther (Seoul) 29, 596–604 (2021). https://doi.org:10.4062/biomolther.2021.122

3 Khoury, S. et al. Quantification of Lipids: Model, Reality, and Compromise. Biomolecules 8 (2018). https://doi.org:10.3390/biom8040174

4 Takats, Z., Wiseman, J. M. & Cooks, R. G. Ambient mass spectrometry using desorption electrospray ionization (DESI): instrumentation, mechanisms and applications in forensics, chemistry, and biology. J Mass Spectrom 40, 1261–1275 (2005). https://doi.org:10.1002/jms.922

5 Pirro, V., Eberlin, L. S., Oliveri, P. & Cooks, R. G. Interactive hyperspectral approach for exploring and interpreting DESI-MS images of cancerous and normal tissue sections. Analyst 137, 2374–2380 (2012). https://doi.org:10.1039/c2an35122f

6 Severiano, D. L. R. et al. Cerebral Lipid Dynamics in Chronic Cerebral Hypoperfusion Model by DESI-MS Imaging. Neuroscience 426, 1–12 (2020). https://doi.org:10.1016/j.neuroscience.2019.11.014

7 Abbassi-Ghadi, N. et al. A Comparison of DESI-MS and LC-MS for the Lipidomic Profiling of Human Cancer Tissue. J Am Soc Mass Spectrom 27, 255–264 (2016). https://doi.org:10.1007/s13361-015-1278-8

8 Czamara, K. et al. Raman spectroscopy of lipids: a review. Journal of Raman Spectroscopy 46, 4–20 (2015). https://doi.org:10.1002/jrs.4607

9 Raman, C. V. & Krishnan, K. S. A New Type of Secondary Radiation. Nature 121, 501–502 (1928). https://doi.org:10.1038/121501c0

10 Orlando, A. et al. A Comprehensive Review on Raman Spectroscopy Applications. Chemosensors 9 (2021). https://doi.org:10.3390/chemosensors9090262

11 Movasaghi, Z., Rehman, S. & Rehman, I. U. Raman Spectroscopy of Biological Tissues. Applied Spectroscopy Reviews 42, 493–541 (2007). https://doi.org:10.1080/05704920701551530

12 Ahlf, D. R., Masyuko, R. N., Hummon, A. B. & Bohn, P. W. Correlated mass spectrometry imaging and confocal Raman microscopy for studies of three-dimensional cell culture sections. Analyst 139, 4578–4585 (2014). https://doi.org:10.1039/c4an00826j

13 Ryabchykov, O., Popp, J. & Bocklitz, T. Fusion of MALDI Spectrometric Imaging and Raman Spectroscopic Data for the Analysis of Biological Samples. Front Chem 6, 257 (2018). https://doi.org:10.3389/fchem.2018.00257

14 Bergholt, M. S. et al. Correlated Heterospectral Lipidomics for Biomolecular Profiling of Remyelination in Multiple Sclerosis. ACS Cent Sci 4, 39–51 (2018). https://doi.org:10.1021/acscentsci.7b00367

15 Noda, I. Two-dimensional correlation and codistribution spectroscopy (2D-COS and 2D-CDS) analyses of planar spectral image data. Journal of Molecular Structure 1211 (2020). https://doi.org:10.1016/j.molstruc.2020.128068

16 Bonifacio, A. et al. Chemical imaging of articular cartilage sections with Raman mapping, employing uni- and multi-variate methods for data analysis. Analyst 135, 3193–3204 (2010). https://doi.org:10.1039/c0an00459f

17 Wang, P. et al. Label-free quantitative imaging of cholesterol in intact tissues by hyperspectral stimulated Raman scattering microscopy. Angew Chem Int Ed Engl 52, 13042–13046 (2013). https://doi.org:10.1002/anie.201306234

18 Guo, S., Rösch, P., Popp, J. & Bocklitz, T. Modified PCA and PLS: Towards a better classification in Raman spectroscopy–based biological applications. Journal of Chemometrics 34 (2020). https://doi.org:10.1002/cem.3202

19 Butler, H. J. et al. Using Raman spectroscopy to characterize biological materials. Nat Protoc 11, 664–687 (2016). https://doi.org:10.1038/nprot.2016.036

20 Kong, K. et al. Diagnosis of tumors during tissue-conserving surgery with integrated autofluorescence and Raman scattering microscopy. Proc Natl Acad Sci U S A 110, 15189–15194 (2013). https://doi.org:10.1073/pnas.1311289110

21 Liu, W., Sun, Z., Chen, J. & Jing, C. Raman Spectroscopy in Colorectal Cancer Diagnostics: Comparison of PCA-LDA and PLS-DA Models. Journal of Spectroscopy 2016, 1–6 (2016). https://doi.org:10.1155/2016/1603609

22 Bergholt, M. S. et al. Raman endoscopy for in vivo differentiation between benign and malignant ulcers in the stomach. Analyst 135, 3162–3168 (2010). https://doi.org:10.1039/c0an00336k

23 Baig, N. F. et al. Multimodal chemical imaging of molecular messengers in emerging Pseudomonas aeruginosa bacterial communities. Analyst 140, 6544–6552 (2015). https://doi.org:10.1039/c5an01149c

24 Petit, V. W. et al. Multimodal spectroscopy combining time-of-flight-secondary ion mass spectrometry, synchrotron-FT-IR, and synchrotron-UV microspectroscopies on the same tissue section. Anal Chem 82, 3963–3968 (2010). https://doi.org:10.1021/ac100581y

25 Bocklitz, T. W. et al. Deeper understanding of biological tissue: quantitative correlation of MALDI-TOF and Raman imaging. Anal Chem 85, 10829–10834 (2013). https://doi.org:10.1021/ac402175c

26 Lanni, E. J. et al. Correlated imaging with C60-SIMS and confocal Raman microscopy: Visualization of cell-scale molecular distributions in bacterial biofilms. Anal Chem 86, 10885–10891 (2014). https://doi.org:10.1021/ac5030914

27 Yan, X. et al. Cell-Type-Specific Metabolic Profiling Achieved by Combining Desorption Electrospray Ionization Mass Spectrometry Imaging and Immunofluorescence Staining. Anal Chem 92, 13281–13289 (2020). https://doi.org:10.1021/acs.analchem.0c02519

28 Wang, Z., Bovik, A. C., Sheikh, H. R. & Simoncelli, E. P. Image quality assessment: from error visibility to structural similarity. IEEE Trans Image Process 13, 600–612 (2004). https://doi.org:10.1109/tip.2003.819861

29 Chan, P. H. Oxygen radicals in focal cerebral ischemia. Brain Pathol 4, 59–65 (1994). https://doi.org:10.1111/j.1750-3639.1994.tb00811.x

30 Bakthavachalam, P. & Shanmugam, P. S. T. Mitochondrial dysfunction - Silent killer in cerebral ischemia. J Neurol Sci 375, 417–423 (2017). https://doi.org:10.1016/j.jns.2017.02.043

31 Alfieri, A. et al. Targeting the Nrf2-Keap1 antioxidant defence pathway for neurovascular protection in stroke. J Physiol 589, 4125–4136 (2011). https://doi.org:10.1113/jphysiol.2011.210294

32 Zhan, Y. et al. Plasma-based proteomics reveals lipid metabolic and immunoregulatory dysregulation in post-stroke depression. Eur Psychiatry 29, 307–315 (2014). https://doi.org:10.1016/j.eurpsy.2014.03.004

33 Lalancette-Hebert, M. et al. Accumulation of dietary docosahexaenoic acid in the brain attenuates acute immune response and development of postischemic neuronal damage. Stroke 42, 2903–2909 (2011). https://doi.org:10.1161/STROKEAHA.111.620856

34 Cai, W. et al. Post-stroke DHA Treatment Protects Against Acute Ischemic Brain Injury by Skewing Macrophage Polarity Toward the M2 Phenotype. Transl Stroke Res 9, 669–680 (2018). https://doi.org:10.1007/s12975-018-0662-7

35 Mohamud Yusuf, A., Hagemann, N. & Hermann, D. M. The Acid Sphingomyelinase/ Ceramide System as Target for Ischemic Stroke Therapies. Neurosignals 27, 32–43 (2019). https://doi.org:10.33594/000000184

36 Gui, Y. K. et al. Plasma levels of ceramides relate to ischemic stroke risk and clinical severity. Brain Res Bull 158, 122–127 (2020). https://doi.org:10.1016/j.brainresbull.2020.03.009

37 Dong, X. et al. Neutrophil Membrane-Derived Nanovesicles Alleviate Inflammation To Protect Mouse Brain Injury from Ischemic Stroke. ACS Nano 13, 1272–1283 (2019). https://doi.org:10.1021/acsnano.8b06572

38 Belayev, L. et al. Docosanoids Promote Neurogenesis and Angiogenesis, Blood-Brain Barrier Integrity, Penumbra Protection, and Neurobehavioral Recovery After Experimental Ischemic Stroke. Mol Neurobiol 55, 7090–7106 (2018). https://doi.org:10.1007/s12035-018-1136-3

39 Bergholt, M. S. et al. Fiberoptic confocal raman spectroscopy for real-time in vivo diagnosis of dysplasia in Barrett’s esophagus. Gastroenterology 146, 27–32 (2014). https://doi.org:10.1053/j.gastro.2013.11.002

40 Heng, H. P. S., Shu, C., Zheng, W., Lin, K. & Huang, Z. Advances in real-time fiber-optic Raman spectroscopy for early cancer diagnosis: Pushing the frontier into clinical endoscopic applications. Translational Biophotonics 3 (2020). https://doi.org:10.1002/tbio.202000018

41 Towers, M. W., Karancsi, T., Jones, E. A., Pringle, S. D. & Claude, E. Optimised Desorption Electrospray Ionisation Mass Spectrometry Imaging (DESI-MSI) for the Analysis of Proteins/Peptides Directly from Tissue Sections on a Travelling Wave Ion Mobility Q-ToF. J Am Soc Mass Spectrom 29, 2456–2466 (2018). https://doi.org:10.1007/s13361-018-2049-0

42 Bergholt, M. S. & Horgan, C. C. in Biomedical Vibrational Spectroscopy 2022: Advances in Research and Industry. PC1195704 (SPIE).

43 Horgan, C. C. et al. High-Throughput Molecular Imaging via Deep-Learning-Enabled Raman Spectroscopy. Anal Chem 93, 15850–15860 (2021). https://doi.org:10.1021/acs.analchem.1c02178

44 Longa, E. Z., Weinstein, P. R., Carlson, S. & Cummins, R. Reversible middle cerebral artery occlusion without craniectomy in rats. Stroke 20, 84–91 (1989). https://doi.org:10.1161/01.str.20.1.84

