## Supplementary information for "Opto-lipidomics of tissues"

**Supplementary Fig. 1**

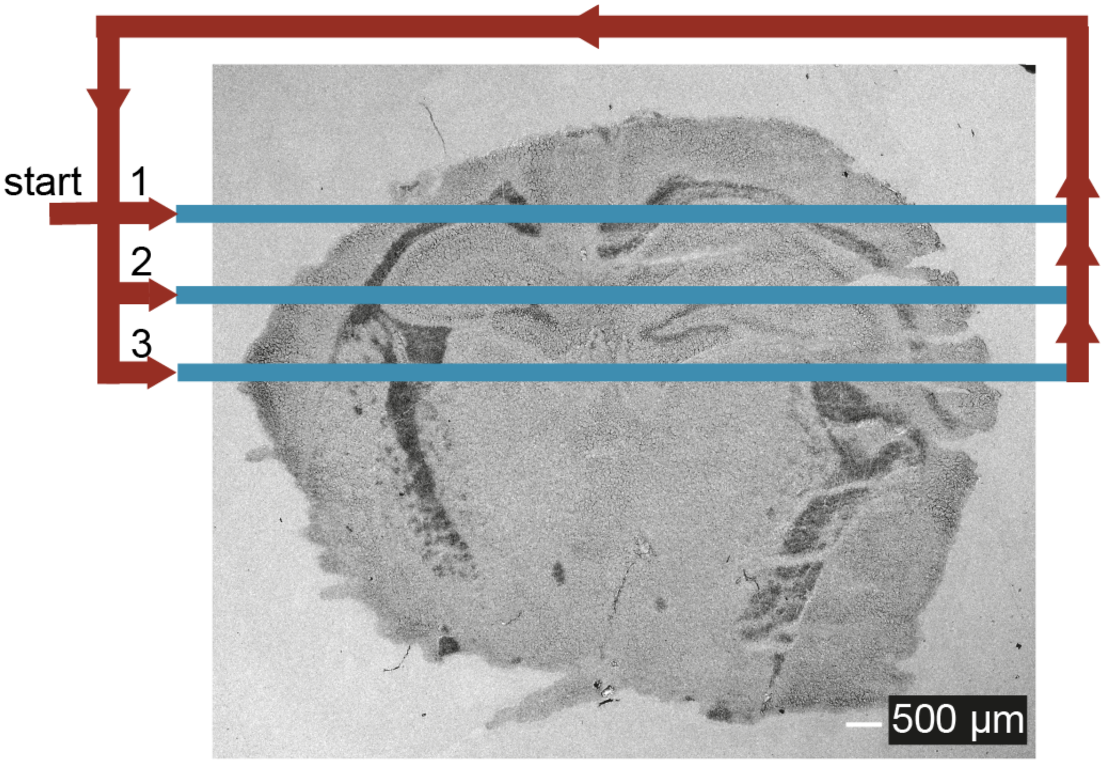


**Supplementary Fig. 1** The scan pattern of the integrated Raman and DESI-MS system. Since the DESI sprayer has a continuous flow of MeOH, the Raman and DESI-MS spectra are acquired only when sampling the tissues. Acquisition stops when the motorised stage moves in a rectangular pattern around the sample to avoid sequential exposure of MeOH.

**Supplementary Fig. 2**

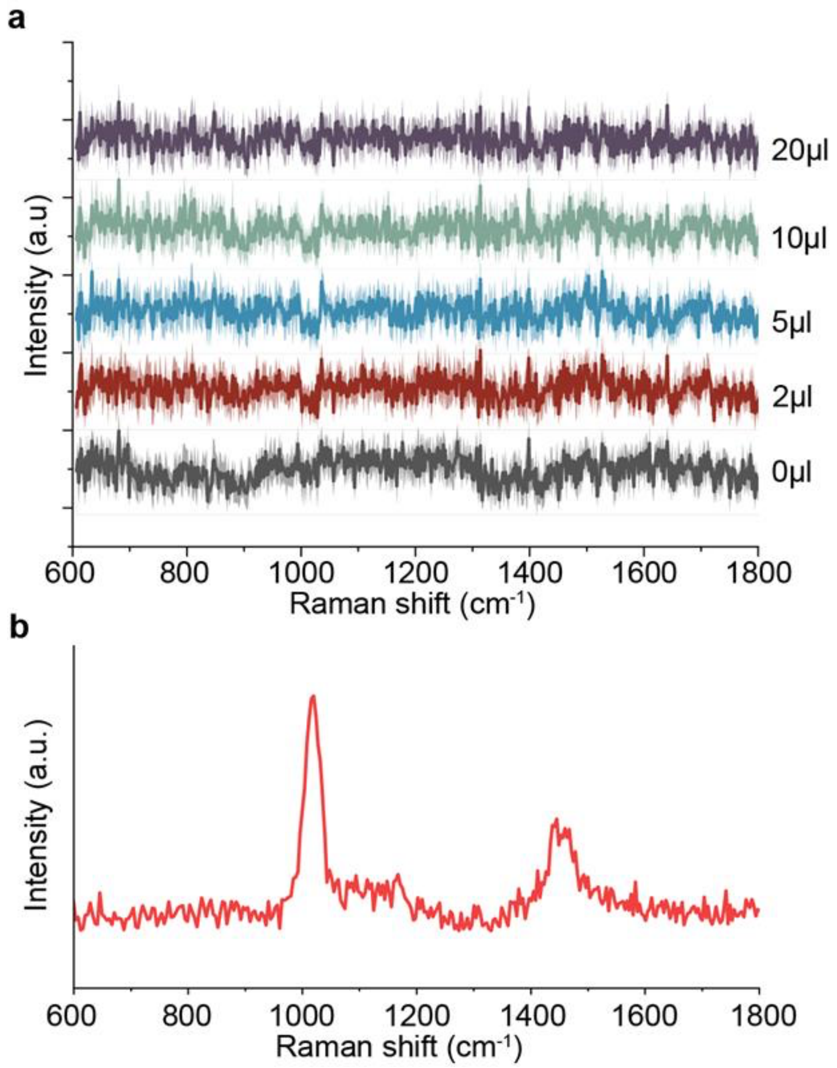


**Supplementary Fig. 2 | a,** Raman spectra ± 1 standard deviation of $MgF_2$ slides with varying methanol flow rates (0-20 µl/min) were collected to investigate the potential contamination of the Raman signal by MeOH peaks. A total of 5 spectra were collected at each flow rate. The integration time for each spectrum was 0.5 seconds. The Raman spectra show no significant contamination by MeOH peaks, even at the highest flow rate of 20 µl/min. **b**, Pure MeOH Raman spectrum showing distinct peaks at near ~1000 $cm^{-1}$ and ~1450 $cm^{-1}$.

**Supplementary Fig. 3**

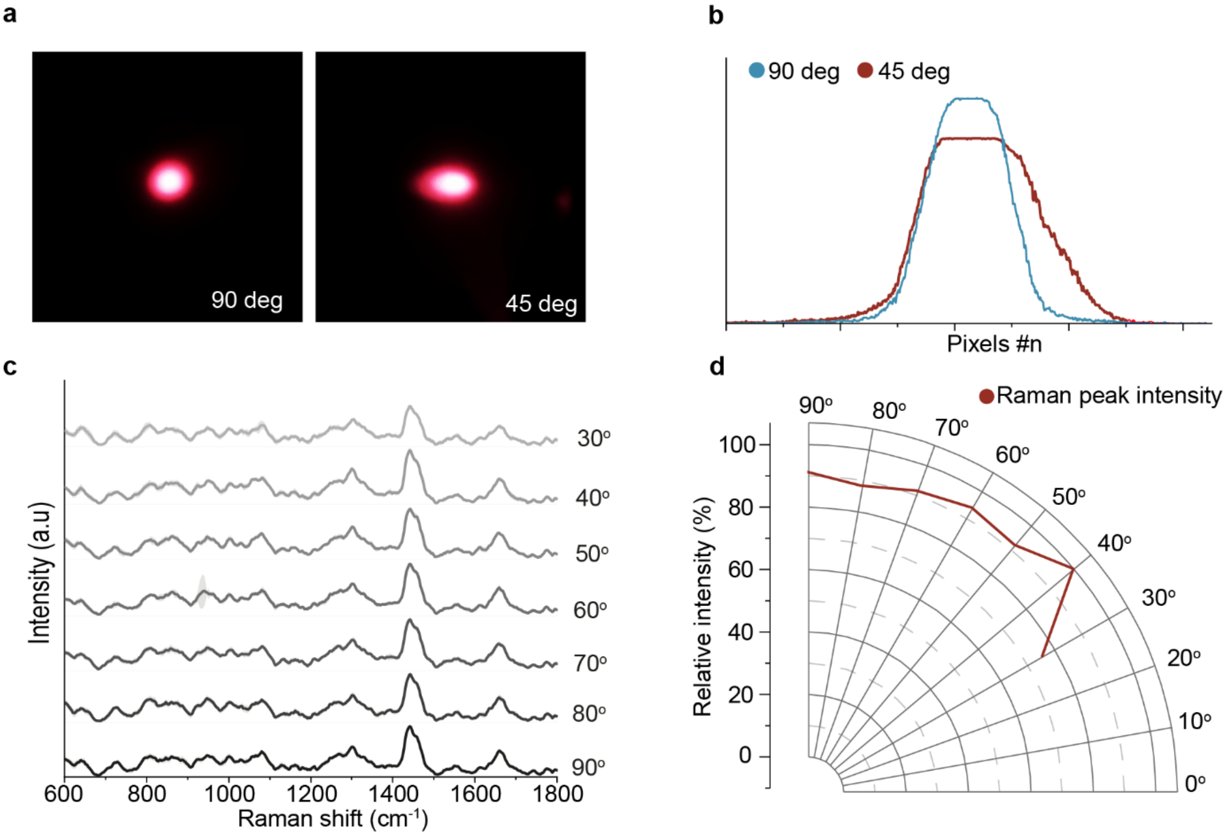


**Supplementary Fig. 3 |** Since the integrated Raman modality collects spectra at an incident angle relative to the surface, we characterised the angular dependence of the tissue Raman spectrum intensity. **a,** Spot shape for the 785 nm laser excitation at 90 degrees and 45 degrees incident angle. **b,** Laser excitation distribution at 90 degrees and 45 degrees incident angle. **c,** Raman spectra of brain tissue at various incident angles 30 - 90 degrees. **d,** Polar plot of the relative intensity of the 1445 $cm^{-1}$ peak at various incident angles 30 - 90 degrees. The tissue Raman signal shows a subtle increase in intensity (~10%) between the normal to the surface and 40 degrees (1445 $cm^{-1}$ peak, $p<0.05$, one-way ANOVA), which is expected for a well-focused spot on a thin tissue section.

**Supplementary Fig. 4**

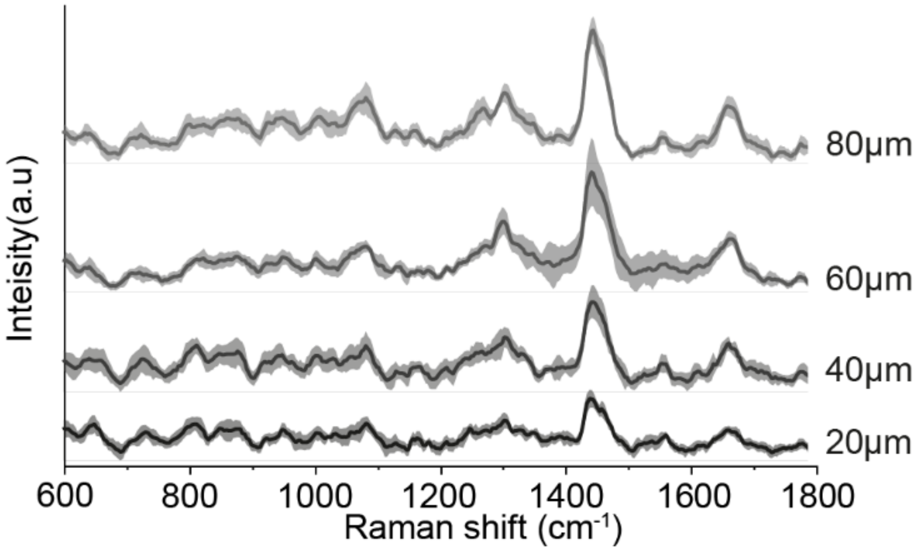


**Supplementary Fig. 4 |** The effect of tissue thickness (20, 40, 60, and 80 μm) on the Raman signal intensity on healthy mouse brain tissue revealed that 40 μm tissue sections provide a good compromise between signal intensity and tissue thickness corresponding to ~2-3x of cell size.

**Supplementary Fig. 5**

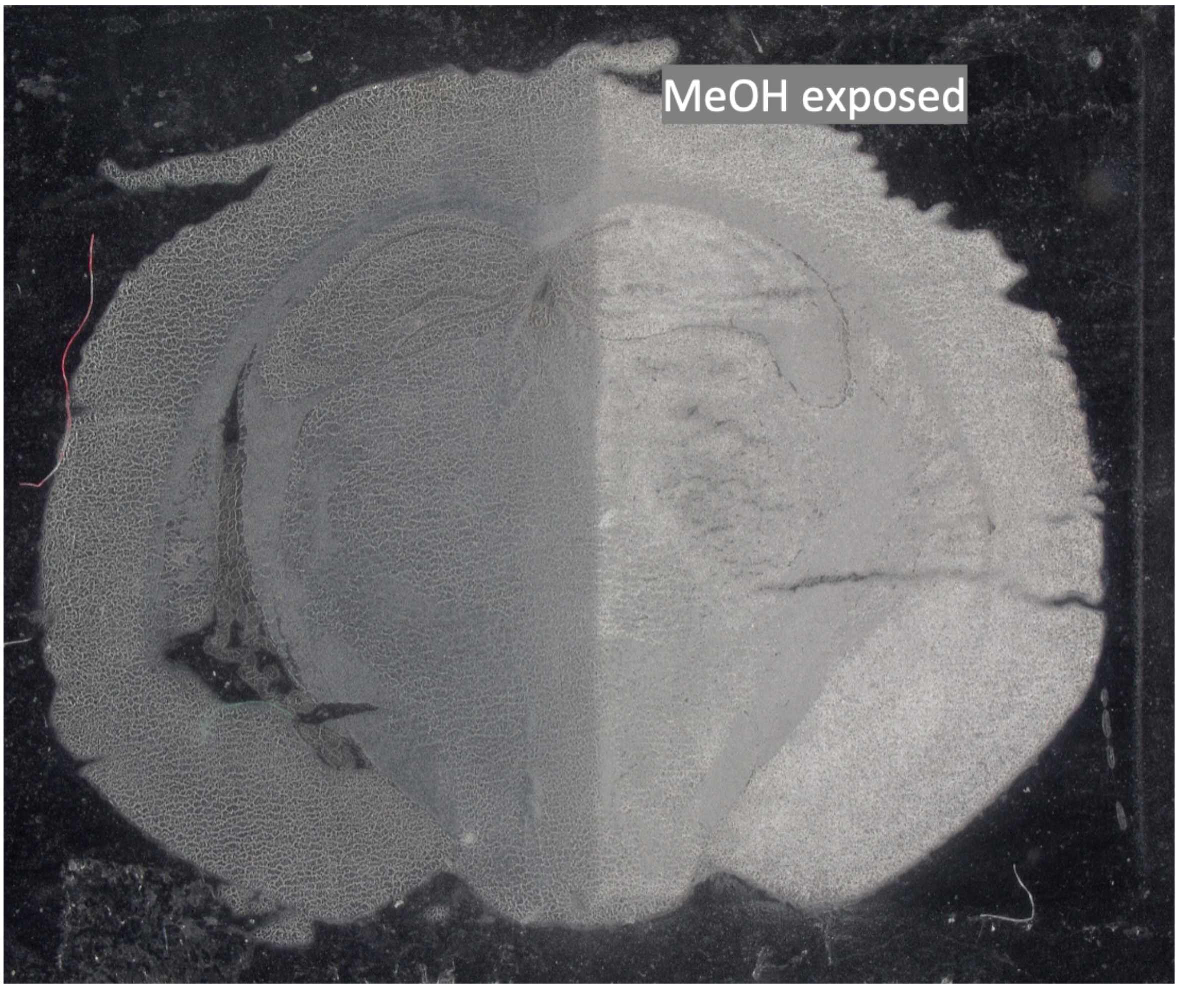


**Supplementary Fig. 5 |** White light image of a mouse brain section illustrates the impact of the DESI-MS measurement on the tissue. The right side of the section has been exposed to MeOH. Visual examination reveals that tissue samples become opaque following DESI-MS imaging.

### Supplementary Fig. 6

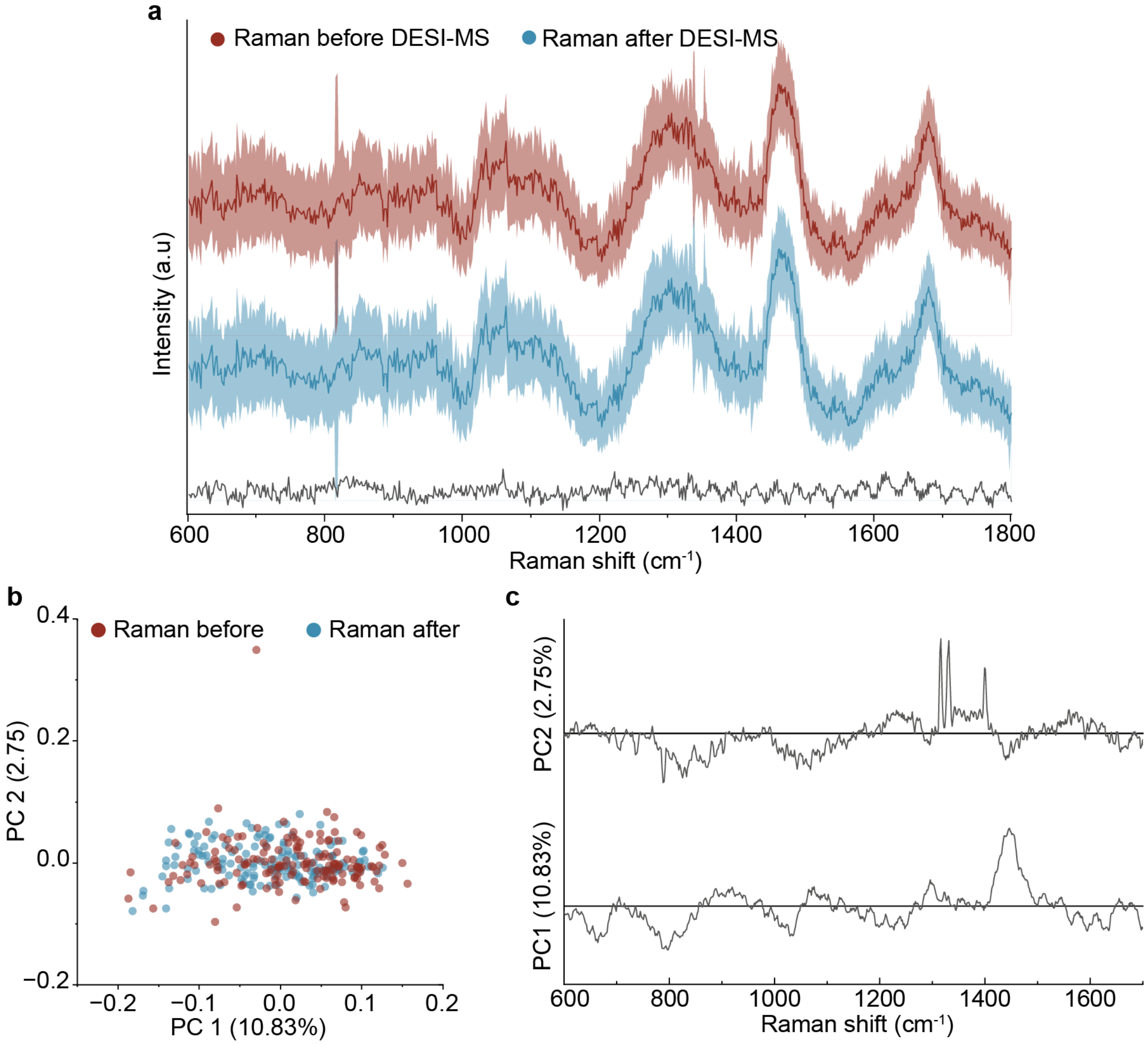


**Supplementary Fig. 6 | a,** Mean Raman spectra of mouse brain tissue before DESI-MS measurement (red), after DESI-MS measurement (blue) (0.5 sec integration time). Also shown is the difference residual spectrum (black). **b,** Principal component analysis (PCA) scores on the Raman data (n=150 before DESI-MS, n=150 after DESI-MS). **c,** PCA loadings reveal noise that were not associated with the DESI-MS imaging.

**Supplementary Fig. 7**

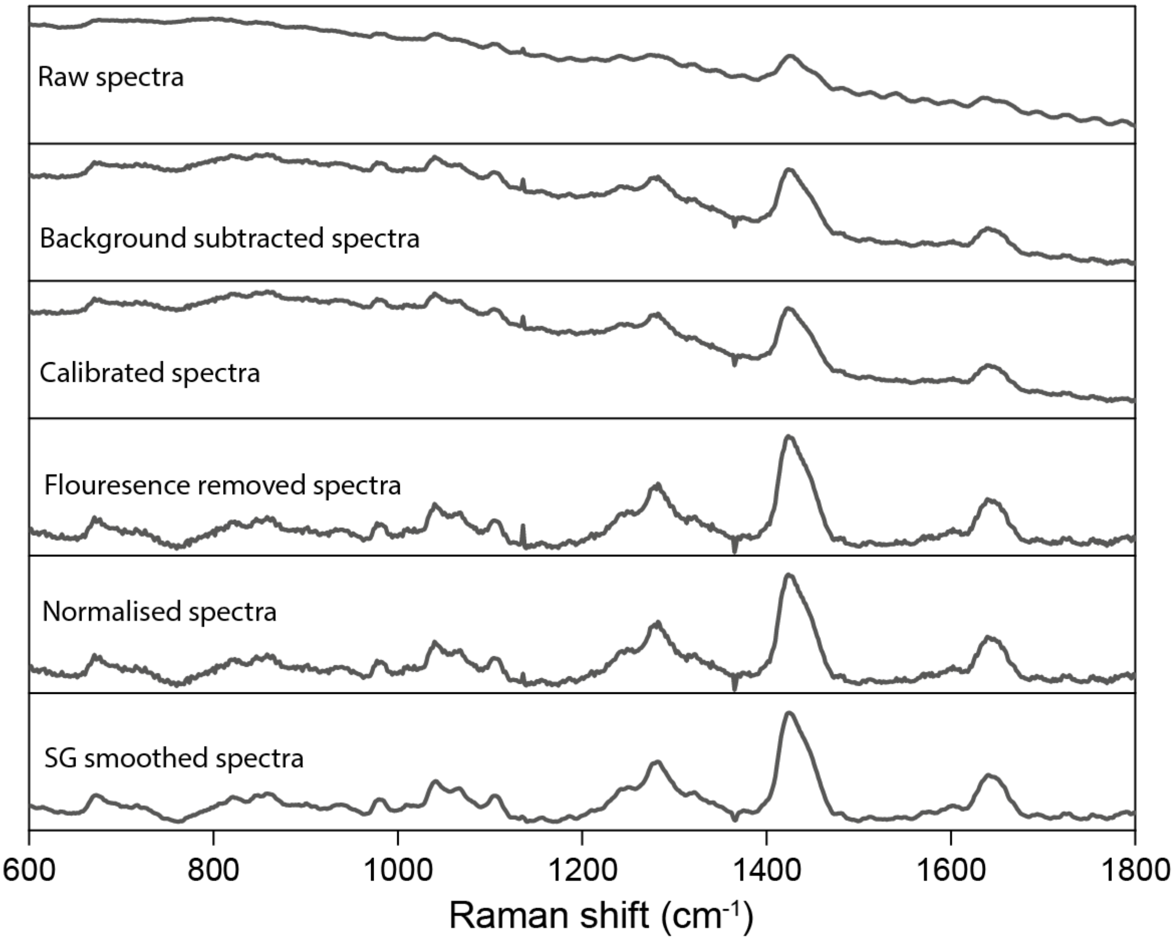


**Supplementary Fig. 7 |** Sequential preprocessing of Raman spectroscopy, including background subtraction, calibration, autofluorescence removal, normalisation and Savitzky Golay (SG) smoothing.

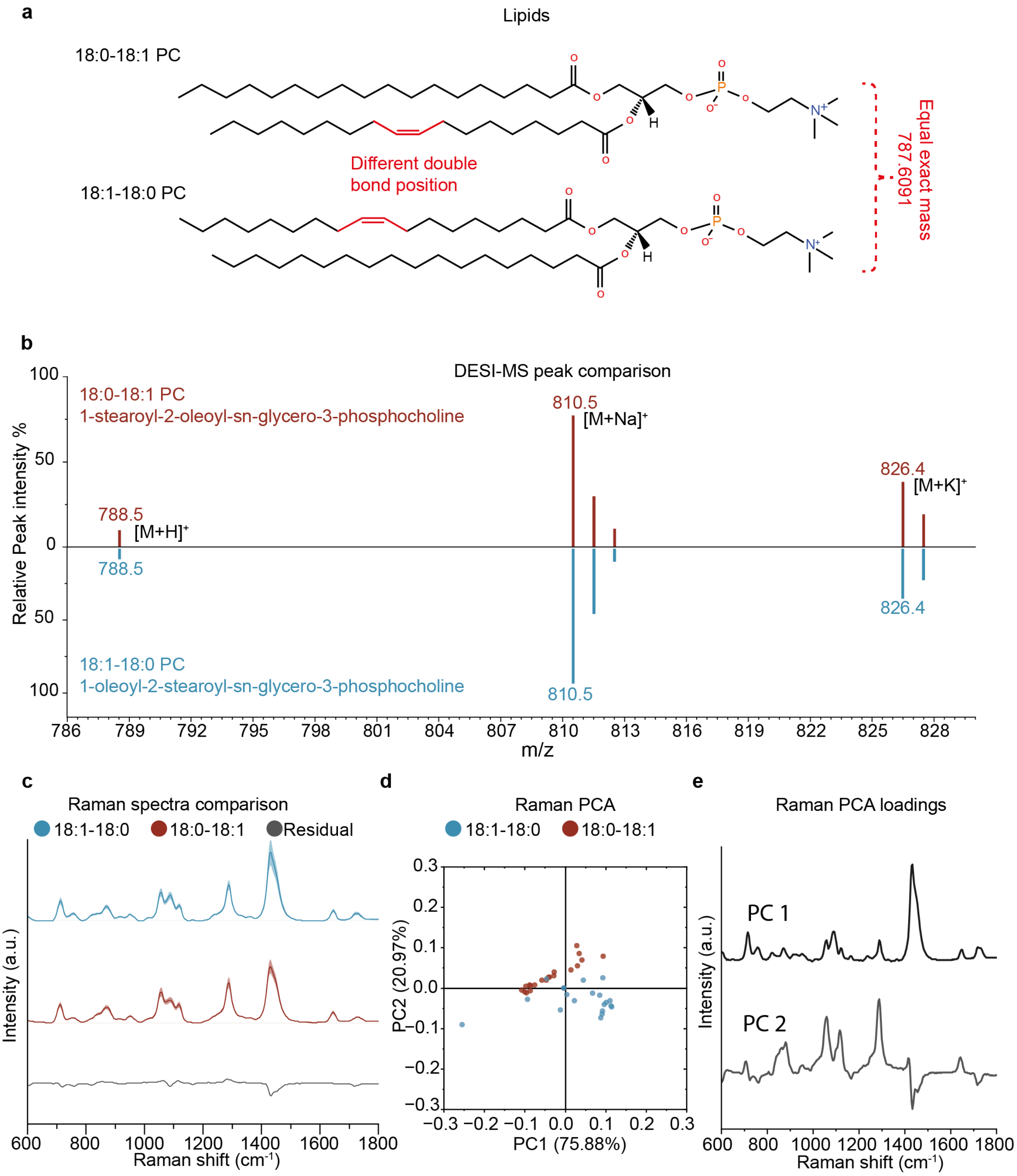


**Supplementary Fig. 8 a,** PC18:1-18:0(1-oleoyl-2-stearoyl-sn-glycero-3-phosphocholine) and PC18:0-18:1(1-stearoyl-2-oleoyl-sn-glycero-3-phosphocholine) of identical elemental composition but with a double band located in a different position of the alkyl chains. **b,** DESI-MS spectrum of the two lipids (mirrored around zero for clarity) showing consistent overlap of the MS peaks. These lipids cannot be discriminated using DESI-MS alone **c,** Average Raman spectra ±1 standard deviation (SD) of the two lipids. Also shown is the difference spectrum (18:1-18:0 minus 18:0-18:1). **d,** Principal component analysis (PCA) scores showing complete separation of the two lipids. **e**. PCA loadings revealing subtle peak shifts in PC2.

**Complementarity of vibrational Raman spectroscopy and DESI-MS**

We established the complementarity between optical spectroscopy and DESI-MS. Species bearing identical elemental composition, such as structural isomers, cannot be differentiated using DESI-MS alone. While tandem MS/MS can often discriminate isomers, the use of Raman spectroscopy could offer a rapid and highly efficient alternative. We investigated if Raman spectroscopy can be used to resolve subtle molecular identity by measuring two very similar lipids: PC18:1-18:0(1-oleoyl-2-stearoyl-sn-glycero-3-phosphocholine) and PC18:0-18:1(1-stearoyl-2-oleoyl-sn-glycero-3-phosphocholine) of equal molecular mass but with an unsaturation positioned differently (Fig. 2a). Positive ion mode DESI-MS spectra showed a complete overlap of the m/z peaks corresponding to fragments the two lipids (Fig. 2b). The average Raman spectra ± 1 SD of the two lipids (Fig. 2c) showed well-known peaks associated with phospholipids, e.g., 1450 $cm^{-1}$ ($CH_2$ deformations), 1650 $cm^{-1}$ υ(C=C) and 1745 $cm^{-1}$ υ(C=O). We also calculated the difference Raman spectrum ± 1 SD revealing subtle differences with consistent spectral peak shifts, particularly at 1066 $cm^{-1}$ and 1426 $cm^{-1}$ corresponding to υ(C-C) and β($CH_2$) bonds of the lipids[9] (Fig. 2c). Two-component PCA analysis and linear discriminant analysis provided 100% discrimination between the two lipid molecules (Fig. 2d-e). This demonstrates that while DESI-MS can offer specific identification of the lipid sub-class, Raman spectroscopy can probe the subtle vibrational differences reflecting the molecular structure and resolve highly similar lipid species. Whilst this experiment was performed with purified lipids and not tissues, it provides an important demonstration of the complementarity of these two techniques. Extracting this information from complex tissues will require more comprehensive correlation with LC-MS/MS.

**Supplementary Fig. 8**

| M/Z acc | M/Z exact | Error (ppm) | Assignment | Elemental formula |
|---|---|---|---|---|
| 283.2653 | 283.2643 | 3.53 | FA 18:0 (Stearic acid) | $C_{18}H_{36}O_2$ |
| 327.2338 | 327.2330 | 2.44 | FA 22:6 (DHA) | $C_{22}H_{32}O_2$ |
| 600.5120 | 600.5128 | -1.33 | Cer 36:1;O2 | $C_{36}H_{71}NO_3$ |
| 788.5416 | 788.5447 | -3.93 | PS 36:1 | $C_{42}H_{80}NO_{10}P$ |
| 834.5290 | 834.5291 | -0.11 | PS 40:6 | $C_{46}H_{78}NO_{10}P$ |
| 885.5504 | 885.5499 | 0.56 | PI 38:4 | $C_{47}H_{83}O_{13}P$ |
| 888.6249 | 888.6240 | 1.01 | SHexCer 42:2;O2 | $C_{48}H_{91}NO_{11}S$ |

**Supplementary Fig. 8 | Table of peaks present in mouse brain experiments.** M/Z acc represents the observed m/z peak value. M/Z exact represents the exact mass of the peak. Error represents the m/z error between the observed and exact m/z in ppm. Assignment represents the assigned species to the peak. Also shown is the elemental formula of the assigned species.

**Supplementary Fig. 9**

| **M/Z exact** | **Assignment** | **Characteristic fragments** | **Observed fragments** |
|---|---|---|---|
| **283.2643** | FA 18:0 (Stearic acid) | 265.2538, 283.2647 | 265.2551, 283.2654 |
| **327.2330** | FA 22:6 (DHA) | 229.1956, 283.2426, 327.2329 | 229.2319, 283.2846, 327.2322 |
| **788.5447** | PS 36:1 | 152.9958, 283.2643, 419.2568, 701.5127, 788.5447 | 152.9967, 283.2653, 419.2616, 701.5201, 788.5463 |
| **834.5291** | PS 40:6 | 152.9958, 283.2643, 419.2568, 747.4970, 834.5291 | 152.9946, 283.2636, 419.2562, 747.4965, 834.5291 |

**Supplementary Fig. 9 | Table of MS/MS fragments of peaks present in mouse brain experiments.** M/Z exact represents the exact mass of the peak. Assignment represents the assigned species to the peak. Also shown are the charateristic fragments of the assigned species as well as the observed fragments observed in the measured MS/MS spectra.

**Supplementary Fig. 10**

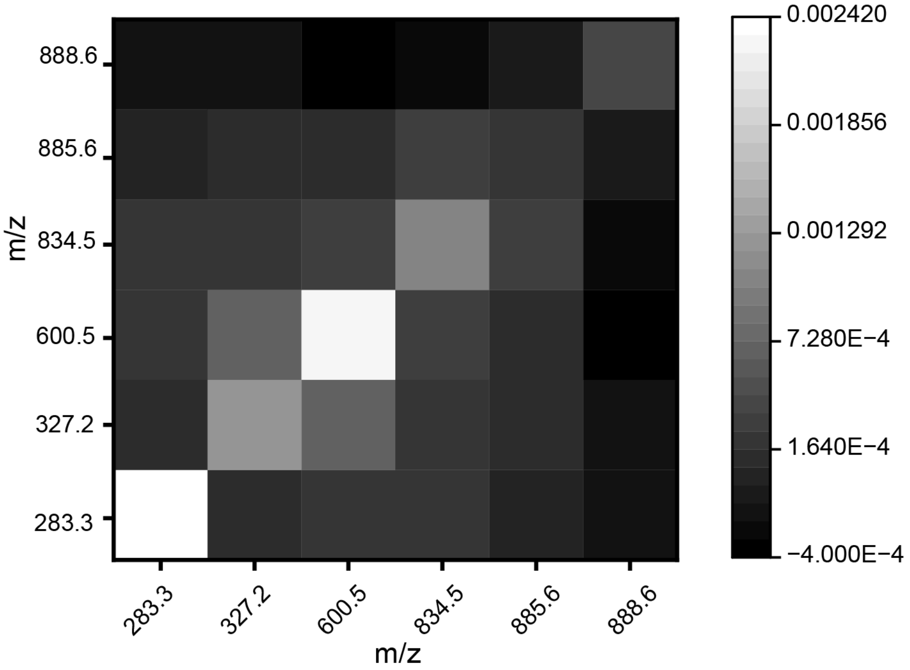


**Supplementary Fig. 10 |** Covariance matrix of some of the major DESI-MS peaks.

**Supplementary Fig. 11**

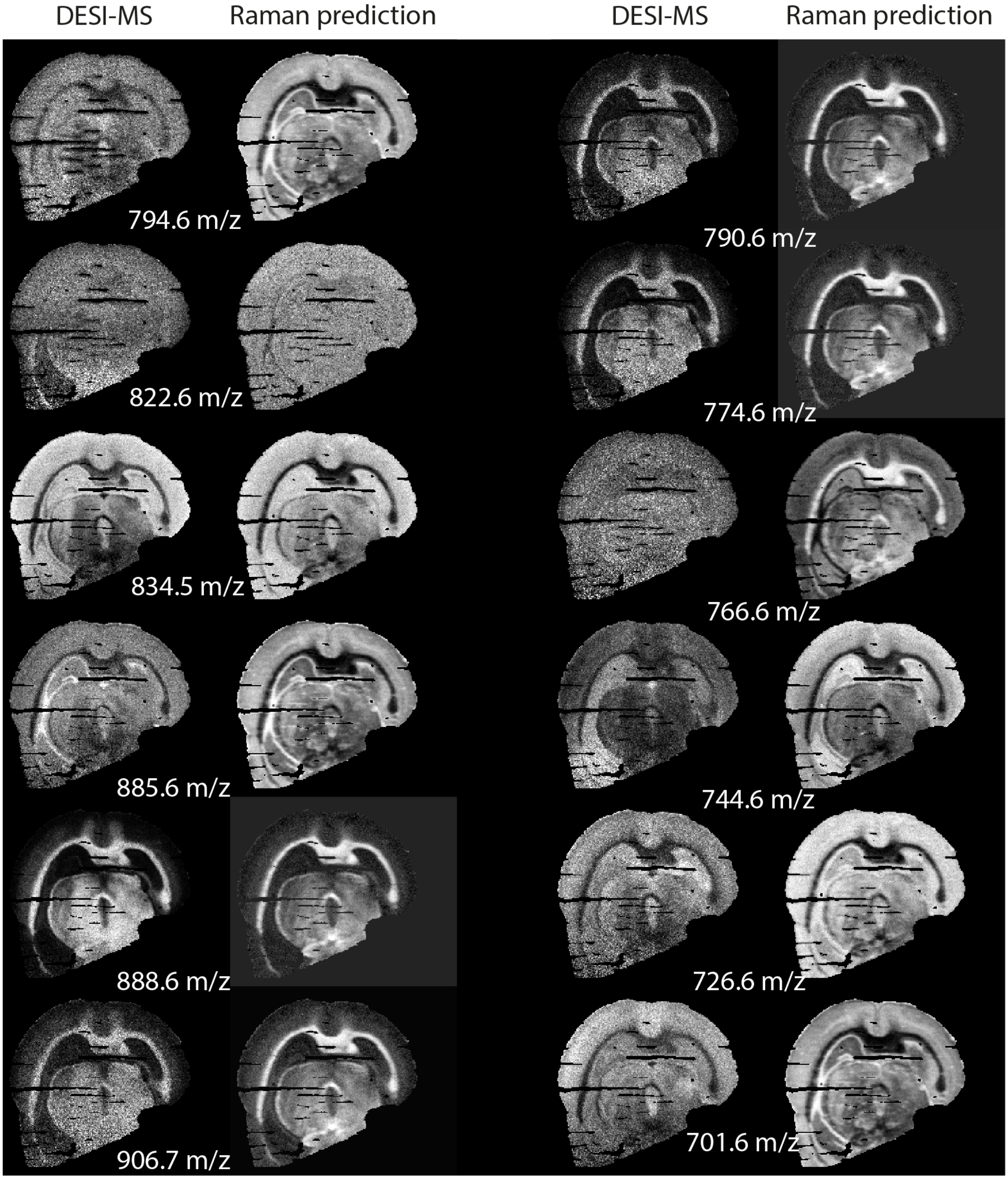


**Supplementary Fig. 11 |** The 12 most intense m/z peak abundance images of DESI-MS. Also shown are the Raman predicted (opto-lipidomics).

**Supplementary Fig. 12**

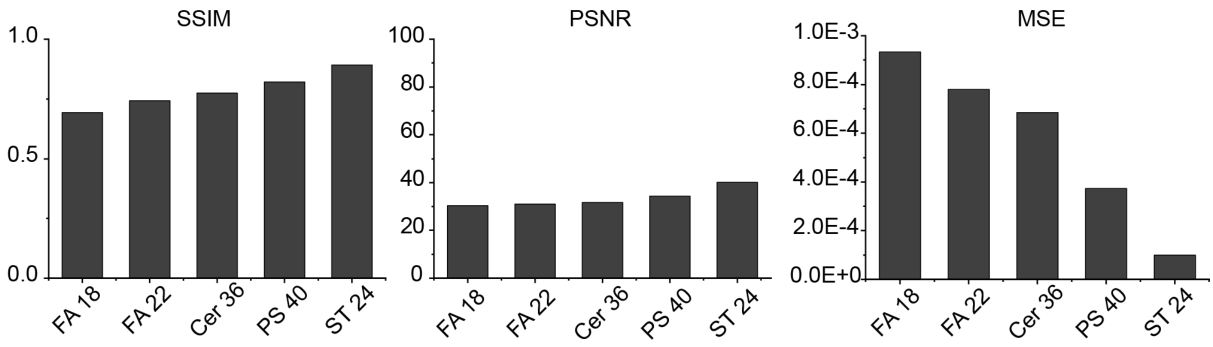


**Supplementary Fig. 12 |** Image quality metrics: **a,** Structural similarity index (SSIM), **b,** Mean squared error (MSE) and **c,** Peak signal to noise ratio (PSNR), for FA 18:0, FA 22:6, Cer 36:1;O2, PS 40:6, and ST 24:1
